# Altered B cells homeostasis in child-onset immunoglobulin A vasculitis

**DOI:** 10.1101/2020.02.28.969444

**Authors:** Deying Liu, Yanfang Jiang, Jinghua Wang, Jinxiang Liu, Meng Xu, Congcong Liu, Sirui Yang

## Abstract

**Background:** Immunoglobulin A vasculitis (IgAV), also called Henoch–Schönlein purpura, is a systemic small vessels vasculitis with immunoglobulin A1-dominant immune deposits. B-cells are a heterogeneous population with unique subsets distinguished by their phenotypes and cytokine production. Here, we explored the status of B cell subsets in patients with IgAV.

**Methods:** Thirty IgAV patients and fifteen age- and sex-matched healthy individuals were enrolled in this study. Fresh blood samples were collected from both healthy and IgAV patients. Upon the distinct expressions of CD3, CD19, CD20, CD38, CD27 and IgD, peripheral blood mononuclear cells (PBMCs) were initially categorized into plasmablasts and memory B cells. Subsequently, using surface markers including CD138 and IgM, and intracellular markers containing IgM and IgG, plasmablasts and memory B cells were further divided into distinct subgroups. A total of eleven populations were detected using multiple flow cytometry.

**Results:** CD3^-^CD19^+^IgD^+^CD27^-^, CD3^-^CD19^+^CD20^-^CD38^+^, CD3^-^CD19^+^CD20^-^CD38^+^IgM^+^, and CD3^-^CD19^+^CD20^-^CD38^+^CD138^+^ B cells were larger in patients with IgAV than in the HCs. Only CD3^-^CD19^+^IgD^-^CD27^+^IgM^+^ B cell counts were reduced in IgAV. The elevated B cell numbers returned to normal after treatment. Plasma and plasmablast B cell numbers correlated with plasma IgA levels. On the contrary, CD3^-^CD19^+^IgD^-^CD27^+^IgM^+^ B cell numbers were negatively proportional to the plasma IgA levels while naïve B cell numbers correlated with plasma and plasmablast B cell counts.

**Conclusions:** We hypothesized that immunoglobulin production was abnormally elevated in IgAV and could be explained by altered B-cell subset homeostasis.

## Background

Immunoglobulin A vasculitis (IgAV), formerly referred to as Henoch–Schönlein purpura, is the most common form of childhood vasculitis. Skin biopsy of the vasculitis lesions reveal small-vessel leukocytoclastic vasculitis [1, 2]. The scarce epidemiological data for childhood IgAV, mainly pertaining to European populations, indicates annual incidence rates from approximately, 3 to 27 in every 100,000 children [3]. Although IgA1-immune deposits, complement factors and neutrophil infiltration in endothelial cells, are widely accepted as characteristic features of IgAV, the casual pathogenic mechanism is yet to be resolved [4, 5]. Additionally, genes also play a crucial role in the pathogenesis of IgAV [6]. It can be triggered by chlamydia, bacteria, viruses, mycoplasma, *Helicobacter pylori*, or infection by parasitic agents. Symptoms include palpable purpura or petechiae, (poly)arthralgia, gastrointestinal disturbances and glomerulonephritis [7]. Usually, IgAV is elf-limiting, but glomerulonephritis in some patients may lead to end-stage renal disease. The prognosis of IgAV is predominantly dependent on the extent of kidney damage [4, 8]. Although IgAV is largely considered as a T cell-mediated disease, several studies have already demonstrated that hyperactivation of T cell subsets, as well as a decline in autoreactive natural killer cell numbers, may also be contributing factors as these cells are key players in the humoral immune response. Additionally, increased serum interleukin (IL)-4, -6, and -17 concentrations have also been found in patients with IgAV [9].

Although T cells have been be involved in human diseases, data on the pathogenesis of B cell subsets are relatively limited. B lymphocytes play a critical role in adaptive immune response, partly by producing high affinity antibodies to pathogens. However, accumulating evidence suggests the pathogenic role of B cells in autoimmune diseases. B cells have a critical role in the initiation and development of several autoimmune diseases such as systemic lupus erythematosus (SLE). IgAV is also caused by destabilized immunity homeostasis. It is thus interesting to explore if pathogenic mechanisms proposed for those diseases may also be involved in IgAV. Upon activation, class switch and differentiation of B cells are regulated by T cells through cytokines and cognate interactions[10]. Additionally, B cells also regulate T cell activation through antigen presentation, production of cytokines and costimulatory molecules, and recruiting T cell subsets and dendritic cells [11]. B-cells are a heterogeneous population with different subsets distinguished by their phenotypes and cytokine production [12]. Several observations regarding the role of plasma B cell in IgAV have been highlighted [13]. It is important to understand the key role of B cells in IgAV, as this may lead to the development of new therapeutic strategies to prevent disease.

However, little information is available on the number of different B cell subsets in patients with IgAV as well as the potential relationship between the subsets. The aim of our study was to describe the altered B cell homeostasis in child-onset IgAV. B cell subsets were determined by flow cytometry using CD19, CD20, CD38, CD138, IgM, and IgG. To the best of our knowledge, we are the first group to show that IgAV patients exhibit an altered peripheral blood B-cell subset distribution.

## Methods

### Patients

Written informed consents were obtained from parents or guardians of all study participants. The experimental protocol followed the guidelines of the Declaration of Helsinki and was approved by the Human Ethics Committee of Jilin University (Jilin University, Changchun, China). Thirty patients were prospectively included if they fulfilled the following criteria : (1) children younger than 18 years of age; (2) met the European League Against Rheumatism/Pediatric Rheumatology International Trials Organization/Pediatric Rheumatology European Society criteria for IgAV [14]: palpable purpura (mandatory) and one of following findings: histopathology (typical LCV with predominant IgA deposits or proliferative glomerulonephritis with predominant IgA deposits); abdominal pain (Diffuse abdominal colicky pain, intussusception and gastrointestinal bleeding); arthritis or arthralgia; renal involvement: proteinuria > 0.3 g/24 h or > 30 mmol/mg of urine albumin/creatinine ratio on a spot morning sample, hematuria [> 5 red blood cells (RBCs)/high-powered field or ≥ 2+ on dipstick or RBC casts in urinary sediment.

Given the self-limiting and benign course of IgAV, symptom-oriented and supportive therapies were administered to patients following admission. Glucocorticoids and/or immunosuppressants (such as cyclophosphamide) were administered. Remission following treatment was defined by two criteria: (1) after 2 weeks, all skin purpura improved, with no appearance of new rashes; and (2) All symptoms were alleviated. Only 25 of the total patients entered remission. There were 3 patients with recurrent skin purpura and 2 with obstinate abdominal pain. We randomly selected 15 patients at the remission stage. The prognosis of IgAV is mostly benign; therefore, blood samples were collected from patients who had successfully entered remission.

A total of 15 sex- and age-matched healthy controls (HCs) were recruited for the study. All participants underwent a routine blood test: measurement of serum immunoglobulin and complement levels using a specific protein analyzer (BN-II; Siemens, München, Germany), serum C-reactive protein (CRP) level using the QuikRead go CRP kit (Orion Diagnostica, Espoo, Finland), urinary protein level using a P800 biochemical analyzer (Roche, Mannheim, Germany), and urinary RBC and white blood cell (WBC) counts using a UF-1000 automatic urinary sediment analyzer (Sysmex, Kobe, Japan).

### Isolation of peripheral blood mononuclear cells

Peripheral blood mononuclear cells (PBMCs) were isolated from HCs and patients with IgAV in the acute and convalescent stages following density-gradient centrifugation using Ficoll-Paque Plus (Amersham Biosciences, Little Chalfont, UK) at 800 × *g* for 30 min at 25°C.

### Flow cytometry

PBMCs at 4 × 10^6^/ml were analyzed by multicolor flow cytometry (FACSAria II; BD Biosciences, Franklin Lakes, NJ, USA). Human PBMCs (10^6^ cells/tube) were stained with CD3, CD19, CD20, CD38, CD138, IgD, CD27, IgM and IgG. We detected 11 subpopulations of B cells: CD3^-^C19^+^, CD3^-^C19^+^CD20^-^CD38^+^ (plasmablasts B cell), CD3^-^C19^+^ CD20^-^ CD38^+^CD138^+^(plasma B cell), CD3^-^C19^+^CD27^-^IgD^+^(naïve B cell), CD3^-^C19^+^CD27^-^IgD^-^ (double negative B cell), CD3^-^C19^+^ CD27^+^IgD^-^(post-switch memory B cell), CD3^-^C19^+^ CD27^+^IgD^-^IgM^+^, CD3^-^C19^+^ CD27^+^IgD^-^IgG^+^,CD3^-^C19^+^CD27^+^IgD^+^ (pre-switch memory B cell)B cells,at room temperature for 30 min. Subsequently, CD3^-^C19^+^CD20^-^CD38^+^ B cells were fixed, permeabilized, and stained with IgG and IgM(a component of the B cell receptor). Data were processed using FlowJo v.5.7.2 software (Tree Star, Ashland, OR, USA).

### Statistical analysis

Data are expressed as the median and range. Kruskal Wallis test was applied to assess the difference among groups. The correlation analysis was evaluated using Spearman’s rank correlation test. The difference between the acute and remission stage was analyzed by the Wilcoxon matched pairs test. Statistical analyses were performed using SPSS 22.0 software. Differences in means were considered statistically significant when two-sided P values were < 0.05.

## Results

### Clinical characteristics

The demographic and clinical characteristics of the study subjects are shown in Table 1. All patients presented with palpable skin purpura, especially on the lower extremities and buttocks. Upon recruitment, the WBC count (P < 0.001), and the platelet (P = 0.0132), serum IgA (P = 0.0243), IgE (P = 0.0411), CRP (P = 0.0213) and complement C4 (P = 0.0467) levels were higher in patients with IgAV than in the HCs (Table 1). No sequelae or other complications were noted.

**Table 1.**
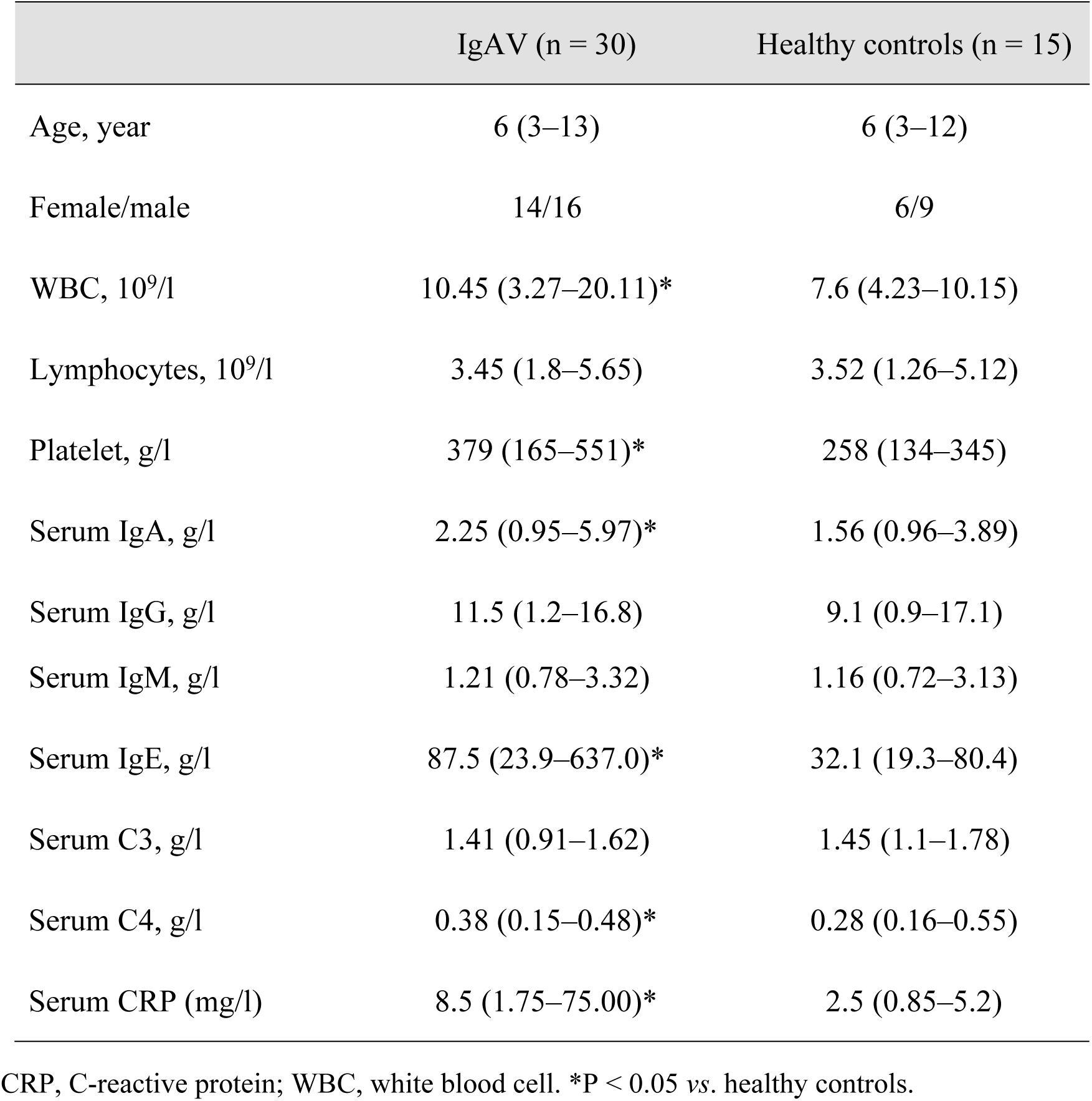
Demographic and clinical characteristics of the study subjects.

|  | IgAV (n = 30) | Healthy controls (n = 15) |
| --- | --- | --- |
| Age, year | 6 (3–13) | 6 (3–12) |
| Female/male | 14/16 | 6/9 |
| WBC, $10^9/l$ | 10.45 (3.27–20.11)* | 7.6 (4.23–10.15) |
| Lymphocytes, $10^9/l$ | 3.45 (1.8–5.65) | 3.52 (1.26–5.12) |
| Platelet, g/l | 379 (165–551)* | 258 (134–345) |
| Serum IgA, g/l | 2.25 (0.95–5.97)* | 1.56 (0.96–3.89) |
| Serum IgG, g/l | 11.5 (1.2–16.8) | 9.1 (0.9–17.1) |
| Serum IgM, g/l | 1.21 (0.78–3.32) | 1.16 (0.72–3.13) |
| Serum IgE, g/l | 87.5 (23.9–637.0)* | 32.1 (19.3–80.4) |
| Serum C3, g/l | 1.41 (0.91–1.62) | 1.45 (1.1–1.78) |
| Serum C4, g/l | 0.38 (0.15–0.48)* | 0.28 (0.16–0.55) |
| Serum CRP (mg/l) | 8.5 (1.75–75.00)* | 2.5 (0.85–5.2) |

CRP, C-reactive protein; WBC, white blood cell. \*P < 0.05 vs. healthy controls.

### Detection of circulating naive and memory B cells in IgAV patients

To investigate the status of B cells in IgAV, we detected the number of four subsets CD3^-^ C19^+^CD27^-^IgD^+^ (naïve B cell), CD3^-^C19^+^CD27^-^IgD^-^ (double negative B cell), CD3^-^C19^+^ CD27^+^IgD^-^ (post-switch memory B cell) and CD3^-^C19^+^CD27^+^IgD^+^ (pre-switch memory B cell) B cells, which were gated from CD3^-^CD19^+^ B cells in flow cytometry analysis of 30 active IgAV patients, 15 patients inremission and HCs (Fig. 1). Circulating CD3^-^C19^+^ and CD3^-^ C19^+^CD27^-^IgD^+^ B cell-counts were increased in active IgAV patients relative to those in the HCs (P = 0.0203, P = 0.0342, respectively) (Fig. 2B). However, the number of circulating CD3^-^ C19^+^CD27^-^IgD^-^, CD3^-^C19^+^ CD27^+^IgD^-^ and CD3^-^C19^+^CD27^+^IgD^+^ cells did not differ between the two groups (P > 0.05; Fig. 2A). Furthermore, we found that CD3^-^C19^+^ CD27^+^IgD^-^ IgM^+^ B cell counts, but not CD3^-^C19^+^ CD27^+^IgD^-^ IgG^+^ B cells were decreased (P = 0.0452; Fig. 2B) in active IgAV compared to that in HCs. All the circulating naive and memory B cells had no difference between the 15 patients in remission and HCs.

**Figure 1.**
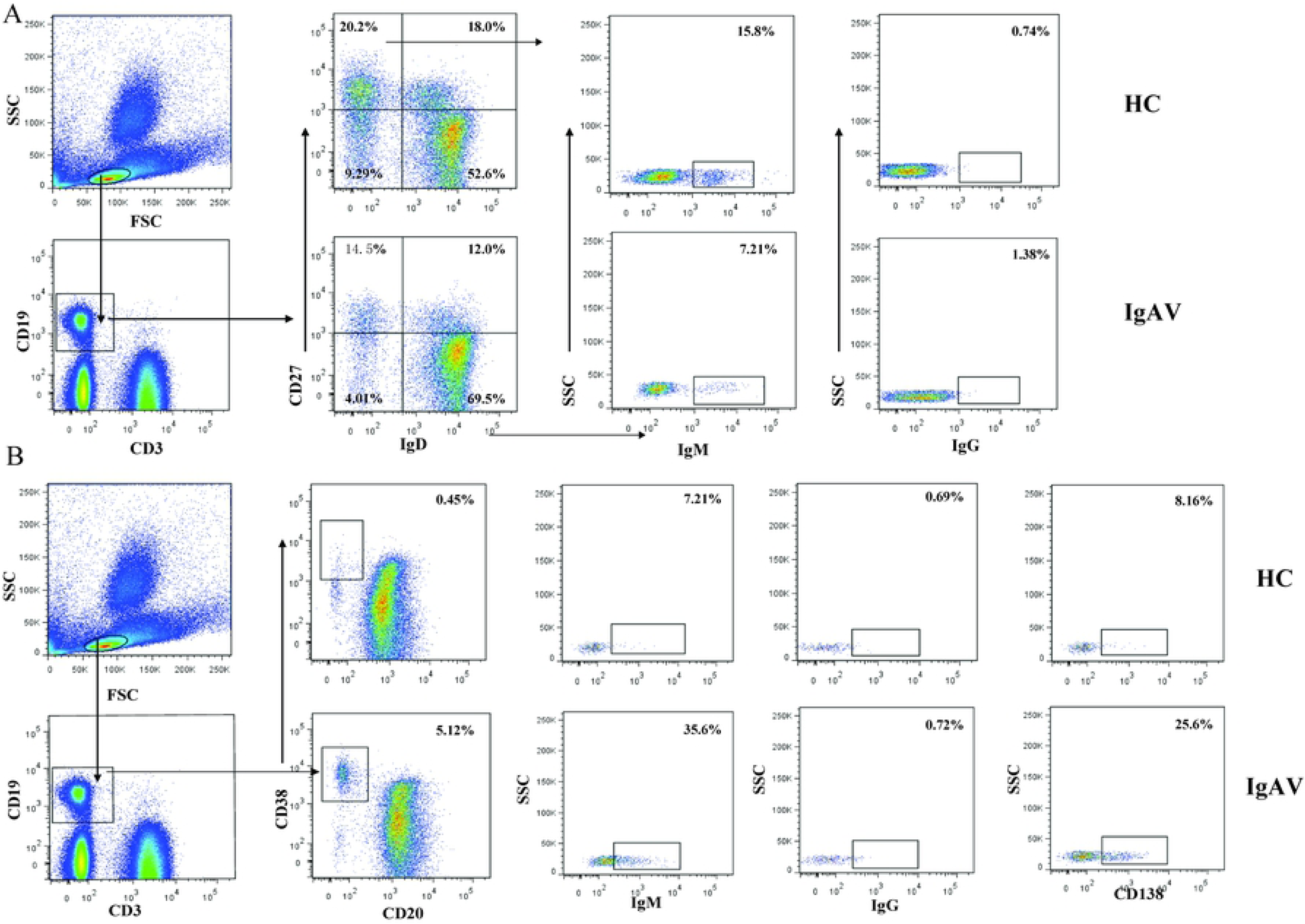
Detection of circulating B cell subsets by flow cytometry. Peripheral blood mononuclear cells (PBMCs) were isolated from patients with immunoglobulin A vasculitis (IgAV) (n = 30) and age and gender-matched healthy controls (HCs; n = 15), labeled with fluorophore-conjugated antibodies, and analyzed by flow cytometry. (A) A gating strategy was used to identify IgD^-^CD27^-^, IgD^+^CD27^-^, IgD^-^CD27^+^, IgD^-^CD27^+^IgM^+^, IgD^-^CD27^+^IgG^+^ and IgD^+^CD27^+^ B cell subsets in CD3^-^CD19^+^ B cell. (B): Gating strategy used to identify CD20^-^CD38^+^, CD20^-^CD38^+^IgM^+^, CD20^-^CD38^+^IgG^+^, CD20^-^CD38^+^CD138^+^ B cell subsets in CD3^-^CD19^+^ B cell.

**Figure 2.**
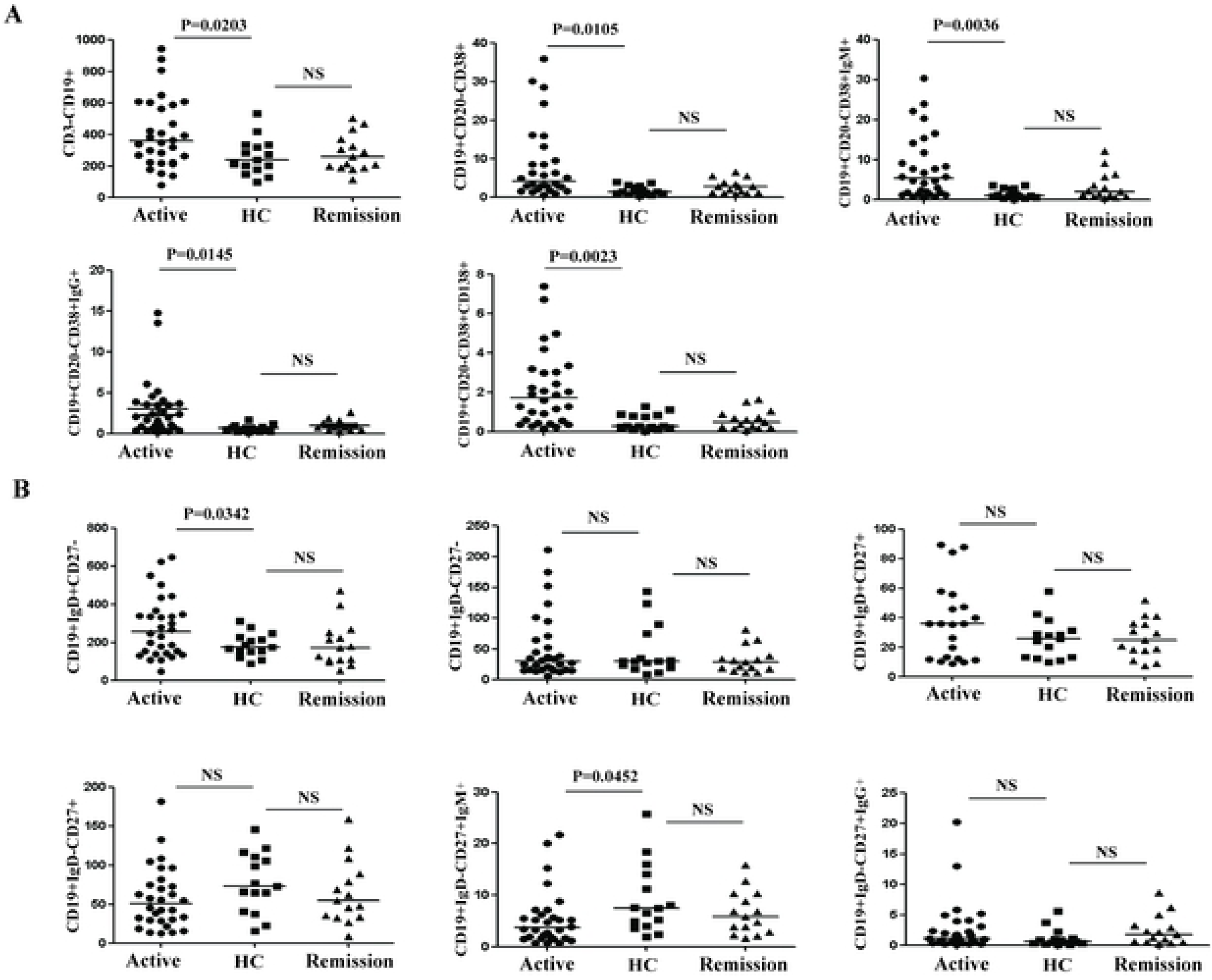
Comparison of different B cell subsets between active and remission in immunoglobulin A vasculitis (IgAV) and HC. NS, not significant.

### Circulating plasmablasts and plasma B cells in IgAV patients

Next, we detected the cell number of four subsets, i.e. CD3^-^C19^+^ CD20^-^CD38^+^ (plasmablasts B cell), CD3^-^C19^+^ CD20^-^CD38^+^CD138^+^ (plasma B cell), CD3^-^C19^+^ CD20^-^CD38^+^IgM^+^, CD3^-^ C19^+^ CD20^-^CD38^+^ IgG^+^ B cells from 30 active patients with IgAV and HCs (Fig. 1B). Circulating CD3^-^C19^+^ CD20^-^CD38^+^ (plasmablasts B cell), CD3^-^C19^+^ CD20^-^CD38^+^CD138^+^ (plasma B cell), CD3^-^C19^+^ CD20^-^CD38^+^IgM^+^ and CD3^-^C19^+^ CD20^-^CD38^+^IgG^+^ B cells in active IgAV were greater than that in HCs (P = 0.0105, P = 0.0023, P = 0.0036 and P = 0.0145, respectively) (Fig. 2A). All the plasmablasts and plasma B cells had no difference between the 15 patients in remission and HCs.

### Alterations in B cell subsets following treatment

Following symptom-oriented and supportive therapies, the majority of patients successfully went into remission. We examined the above B cell subsets in 15 patients in remission (Fig. 3). Naïve, plasmablasts, plasma B cells were reduced relative to the absolute value in the active stage (P = 0.0383, 0.0026, and 0.0020, respectively). There were no changes in CD3^-^ C19^+^CD27^+^IgD^-^ (P > 0.05; Fig. 3C) and CD3^-^C19^+^CD27^+^IgD^-^IgM^+^ (P > 0.05; Fig. 3E) levels during remission.

**Figure 3.**
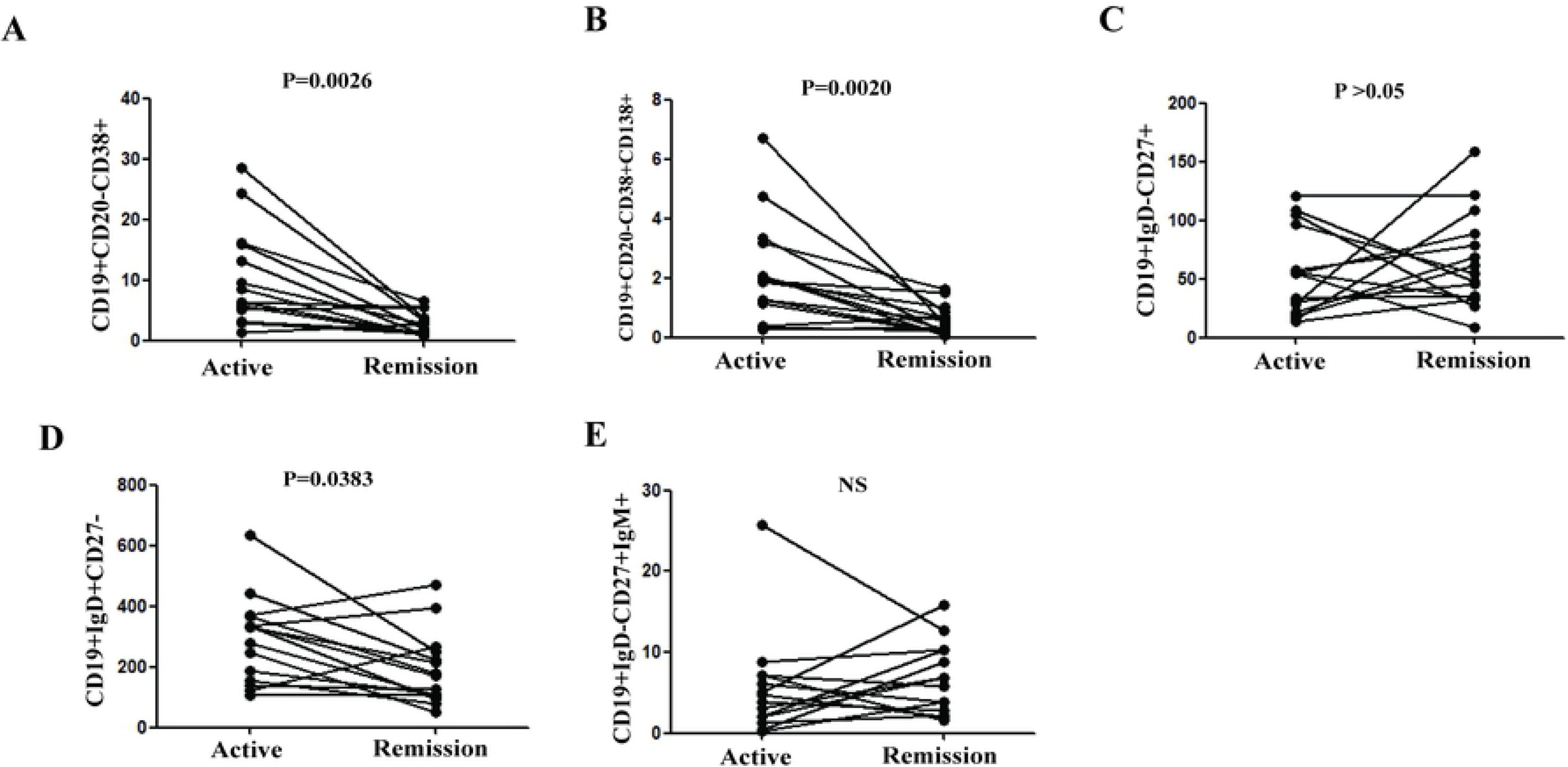
Treatment-induced changes in B cell subsets. Following treatment, 15 patients exhibited remission from disease. B cell counts were compared between active and remission stages.

### Association between B cell subsets and clinical parameters

We investigated whether alterations in B cell subsets were associated with disease etiology and progression, and found that the number of CD19^+^CD20^-^CD38^+^ (r = 0.4320, P = 0.0171), CD19^+^CD20^-^CD38^+^CD138^+^ (r = 0.5316, P = 0.0025),CD19^+^CD20^-^CD38^+^IgM^+^ (r = 0.5847, P = 0.0007), but not of CD20^-^IgD^+^CD27^-^ (r = 0.1393, P = 0.4628) B cells was positively correlated with serum IgA levels (Fig. 4A). By contrast, circulating CD19^+^IgD^-^CD27^+^IgM^+^ B cell counts were inversely related to serum IgA levels (r = −0.3755, P = 0.0409; Fig. 4A). Additionally, CD19^+^CD20^-^CD38^+^ (r = 0.0545, P = 0.7749), CD19^+^CD20^-^CD38^+^CD138^+^ (r = −0.1008, P = 0.5962), CD19^+^CD20^-^CD38^+^IgM^+^ (r = 0.1174, P = 0.5369),CD19^+^IgD^+^CD27^-^ (r = 0.0520,P = 0.7850) and CD19^+^IgD^+^CD27^-^IgM^+^ (r = −0.0092,P = 0.6019) cell subsets showed no association with serum C4 in the IgAV, respectively. We also explored the relationship between B cell subsets and serum IgA level/C4 in the HCs, there is no statistical significance (data not shown).

**Figure 4.**
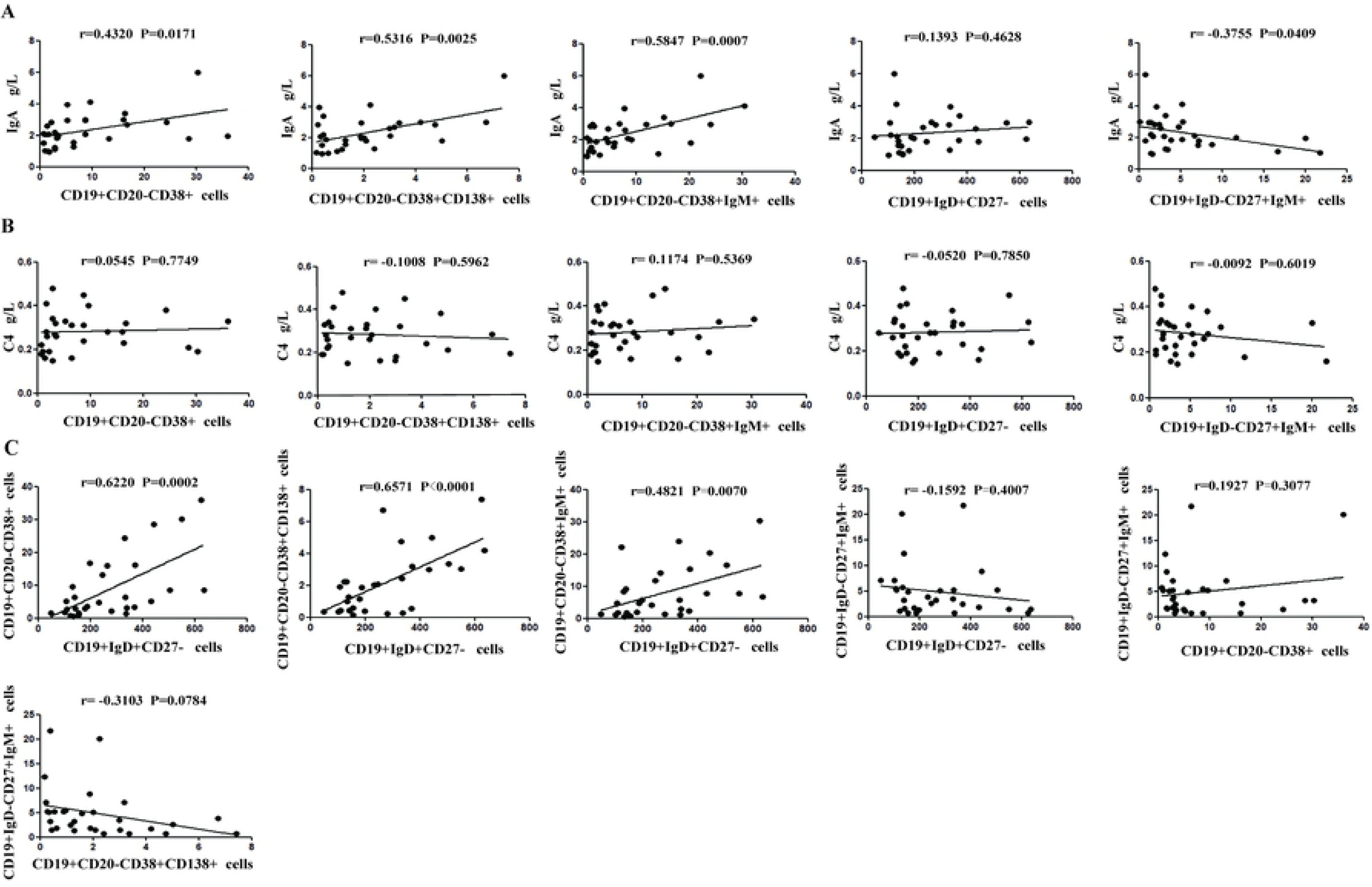
Correlation between and among B cell subsets and serum IgA or plasma C4 levels. Correlations between indicated B cell subsets and (A) serum IgA level, (B) plasma C4, (C) among B cell subsets were analyzed by Spearman’s rank correlation test.

### The relationship among the different B cell subsets

Meanwhile, we analyzed the potential relationship between the numbers of the circulating B cell subsets in the IgAV patients. Circulating CD19^+^IgD^+^CD27^-^ naïve B cell counts correlated with the number of CD19^+^CD20^-^CD38^+^ (r = 0.6620, P = 0.0002), CD19^+^CD20^-^CD38^+^CD138^+^ (r = 0.6571, P < 0.0001), CD19^+^CD20^-^CD38^+^IgM^+^ (r = 0.4821, P = 0.0070) B cells. But there was no relationship observed between CD19^+^CD20^-^CD38^+^ (r = 0.1927, P = 0.3077), CD19^+^CD20^-^CD38^+^CD138^+^ (r = −0.3103, P = 0.0784), CD19^+^IgD^+^CD27^-^(r = 0.1592, P = 0.4007) and CD19^+^IgD^-^CD27^+^IgM^+^ B cells.

## Discussion

IgAV is a common vasculitis with an early age of onset, triggered by environmental and genetic factors, and is associated with a history of often URTI. Although the exact pathogenesis has not yet been elucidated, interaction between the T and B cell lineages is considered a key underlying cause. It has long been presumed that aberrant deposition of glycosylated IgA_1_ and complement activation contribute to IgAV. In this study, increased levels of IgA were found in the patient’s peripheral blood, IgA is mostly known as the major antibody subset present in mucosal areas, where it plays a key role in mucosal defense. Approximately 90% of the IgA present in the circulation is IgA1 while <10% is IgA2. It is thus plausible that the level of IgA in peripheral blood is equivalent to IgA1. When immune complexes are deposited, they activate the complement pathway. Complement is present in inactive form in the circulation, and three pathways can lead to the activation of complement. However, it has been demonstrated that IgA can induce the mannan-binding lectin and alternative complement pathways. GalNAc on the surface of pathogens may facilitate the production of cross-reactive IgA and IgG, which recognize Gd-IgA1 (Galactose-deficient IgA1) [15]. We believe that Gd-IgA1 complexes are deposited in the vasculature due to the aberrant elevation of B cell numbers, thereby inducing neutrophil migration and activation with concomitant tissue damage. Vascular damage is induced by IgA via inflammatory processes including antibody-dependent cell-mediated cytotoxicity (ADCC), reactive oxygen species (ROS) production, and neutrophil extracellular traps (NETs) formation. Additionally, IgA stimulation of neutrophils leads to the release of LTB4, inducing subsequent neutrophil migration in a positive feedback loop [15].

In our previous work, we demonstrated that Tfh cell subpopulations contributed differentially to IgAV pathogenesis and remission [9]. T follicular helper (TFH) cells are specialist providers of B cell help, notably by the secretion of IL-21. Upon activation, B cells interact with TFH cells and can promote the maturation of B cell response within GCs, thereby leading to autoantibody secretion. Although not completely understood, the situation is somewhat similar in the B cell compartment. In fact, the expansion of CD19^+^ B cells has been implicated in IgAV by a previous study [13]. In support of this data, we detected a greater level of B cell subsets in IgAV and believe that an altered balance of circulating B cell subsets may be associated with IgAV.

B cells, as part of the adaptive immunity, are responsible for the humoral responses against pathogens and produce a significant amount of antibodies [16]. Dysregulated frequency of T and B cells can be found in various human autoimmune diseases such as rheumatoid arthritis, systemic lupus erythematosus, and multiple sclerosis, with different cytokines, transcription factors. From studies conducted previously in our lab, we learned that IgAV is mainly mediated by T helper type 2 cell immunity. In this study, we show a slew of evidence demonstrating the involvement of B cells and their antibody production in the IgAV course.

B cells contribute to disease pathogenesis in autoimmunity by presenting antigens as well as stimulation via cytokines to T cells [17-21]. B cells also play an immunomodulatory role in regulating the immune response by secreting cytokines that inhibit disease onset and/or progression [10, 22]. Recent progresses in B-cell activation and differentiation suggested a complex scenario with multiple steps in the generation of long-lived plasma cells (PCs) and memory B cells in the follicles of germinal centers (GCs) as well as in extra-follicular plasmablasts [16, 23-25]. B cell activation is triggered by antigen recognition through B-cell receptor (BCR) either directly or with the help of antigen presenting cells (APCs) in peripheral lymphoid organs, and is achieved by the activation of intracellular signaling pathways and subsequent target gene expression [23]. The activated B cells migrate to the B-T area of lymphoid organs where they undergo a limited expansion upon cognate interaction with antigen-primed T cells. A fraction of memory B cells differentiate into short-lived plasmablasts providing prompt responses to antigens, while others initiate the formation of GC in secondary follicles. Notably, we also observed that IgAV patients have significantly higher levels of peripheral blood CD19^+^CD20^-^CD38^+^ and CD19^+^CD20^-^CD38^+^CD138^+^ B cells when compared to healthy individuals. However, after treatment these B cell subpopulations return to normal levels suggesting their participation in the pathogenic process of IgAV and that they may be used for monitoring disease progression. At the same time, the abnormal increase in naïve B cells can also promote the production of plasma B cells. With strong correlation between peripheral CD19^+^CD38^+^, CD19^+^CD38^+^CD138^+^ B cells, and total IgA levels in IgAV patients. We concluded that plasma B cells are a distinct B cell subset contributing to abnormal IgA production in IgAV. Our results highlight the association between systemic immunity and IgAV pathogenesis. Furthermore, treatments resulted in significant decrease in the proportion of peripheral plasmablasts. The CD19^+^CD38^+^, CD19^+^CD38^+^CD138^+^ B cell subsets may facilitate the production of IgA. T-B cell interactions can increase the production of IgA which aggravates vascular damage and that in turn further activates the immune system. Thus, all the cells and cytokines get involved in a vicious circle of immune activation.

Although the exact function of all the B cell subset is not entirely understood, it is known that it belongs to the memory B cell compartment because of the high levels of somatic hypermutation. Moreover, it has been suggested that these cells might contribute to inflammation by induction of T cell responses and the production of proinflammatory cytokines. It is possible that these B cells are linked with the production of higher affinity antibodies relevant in inflammation. Indeed, subsets of memory B cell shave been reported to dysregulated in other autoimmune diseases such as systemic lupus erythematosus and multiple sclerosis[18]. Based on the reported pathogenic role of B cell in other autoimmune diseases[11], we anticipated that also plays an important role in etiology of IgAV. In the present study, the frequency of IgD^+^CD27^-^ naïve B cells were higher in IgAV patients than controls but have no correlation with the serum level of IgA. Alternatively, naïve B cells may not play a major role in the pathogenesis of IgAV. Nevertheless, naïve B cell numbers as well as the level of plasma IgA correlated with CD19^+^CD20^-^CD38^+^, CD19^+^CD20^-^CD38^+^CD138^+^ and CD19^+^CD20^-^ CD38^+^IgM^+^ B cells. Therefore, we speculate that naïve B cell subset is indirectly involved in the pathogenesis of IgAV by promoting the generation of plasmablasts B cells. We characterized the numbers of circulating CD19^+^IgD^-^CD27^+^, CD19^+^IgD^-^CD27^-^, CD19^+^IgD^+^CD27^+^ B cells and found no significant difference between IgAV patients and HCs. It is possible that antigen may activate memory B cells, which differentiate into plasma cells, leading to antigen-specific IgA production and the pathogenesis of IgAV. We observed a significant reduction in the number of CD19^+^IgD^-^ CD27^+^IgM^+^ memory B cells. Furthermore, CD19^+^IgD^-^CD27^+^IgM^+^ memory B cells is negatively correlated with the level of plasma IgA. Classically, the IgD-CD27+ memory B cells have switched their IgM to IgG, IgA or IgE, IgM^+^ memory B cells maybe the frontline responders by directly giving rise to IgM-secreting cells. It is possible that antigen may activate memory B cells, which differentiate into plasma cells, leading to antigen-specific immunoglobulin production and the pathogenesis of IgAV. Therefore, CD19^+^IgD^-^CD27^+^IgM^+^ memory B cells are exhausted and may be a sensitive marker for evaluating plasma cells. Collectively, our data suggest that the decreased numbers of this distinct group of memory B cells may be associated with the development of IgAV.

Abnormal B-cell activation and differentiation in antibody-driven autoimmune diseases is one of the hallmarks with the continuous production of autoantibodies [23]. B cells can participate in the pathogenesis of autoimmune diseases via several mechanisms: as cytokine producers, antigen-presenting cells, or autoantibody secretors. Elevated IgA is one of the key features of IgAV but it is unclear how B cell activated in IgAV. We suspected that these B cell numbers increased via the activation of signaling mediated through BCR and/or its associated molecules. We and others have reported that a combination of BCR-triggering, costimulatory signals including either CD40L, soluble BAFF or IL-21 and Toll-like receptors (TLRs) stimulation induces the most robust B cell activation and differentiation. B cell activating factor (BAFF, also known as TNF ligand superfamily member 13B) is a key cytokine that promotes the maturation, proliferation and survival of B cells [26]. BAFF has been suspected to play a role in progressive systemic sclerosis (pSS) on the basis of elevated BAFF levels in the serum of patients with pSS and the correlation of BAFF levels with levels of antiSSA/Ro and anti-SSB/La antibodies and RF [27]. BAFF could provide a link between activation of the innate immune system and the adaptive immune system (mainly via B cell stimulation. We speculate that BAFF may be associated with IgAV.

Although IgAV is benign and self-limiting, and majority of patients enter remission, the degree of renal damage determines the long-term prognosis of the IgAV patients, particularly males > 10 years of age with severe gastrointestinal symptoms (abdominal pain, gastrointestinal bleeding, and severe bowel angina), arthritis/arthralgia, persistent purpura or relapse, WBC > 15 × 10^9^/L, platelets > 500 × 10^9^/L, elevated ASO, and low C3 [28]. Henoch-Schönlein purpura nephritis (HSPN) is a major cause of mortality and morbidity in children with HSP, which occurs in 30%-80% of patients during the first three weeks of the initial presentation [29]. For the refractory and severe IgAVN patients, the unraveling of the exact pathogenesis of IgAV will provide directions for prevention of disease, identification of biomarkers and future therapeutic remedies. B cell–targeted approaches for treating immune diseases of the kidney and other organs have gained significant momentum [10, 30]. With our study demonstrating that IgAV patients have abnormal circulation of B cell subsets, B cells gain prominence as the causative agents in IgAV pathogenesis and the over-activation of B cells is the result of a multistep process. Environmental triggers occurring in the presence of genetic and epigenetic dysregulation of IgAV patients lead to the stimulation of specific B cell subsets, particularly plasmablasts and plasma cells. For this reason, it has been considered as a candidate therapeutic target. This is reinforced by the therapeutic efficacy of rituximab, an anti-CD20 monoclonal antibody that specifically depletes B cells. Understanding its function and signaling mechanisms would provide more tools to look for additional therapeutic targets. However, it is well known that IgAV is clinically heterogeneous which leads to failure in developing efficacious targeted therapies. Some studies suggest that rituximab is an effective and safe therapeutic option for adult-onset IgAV [31]. Similarly, B cell–depleting therapy may also be an alternative treatment for patients with IgAN or IgAVN and nephritic–nephrotic syndrome[32]. Identification of the multiple factors that support B cell activation has led to the development of promising targeted therapies, especially for the intractable patients.

Our study had some limitations such as a relatively small sample size and the lack of functional study of memory B cells and plasmablasts in the pathogenic process of IgAV. We are also interested in further investigating the values of these subsets of cells in the kidney lesion to understand their roles in the pathogenesis of IgAV. Biomarkers, associated with the risk of nephritis and renal sequelae, must also be investigated in future.

### Conclusions

In summary, we show that abnormal distribution of B cell subsets in IgAV patients may play a causal role in development of the disease and should be targeted by future therapeutic efforts.

## Declarations

## Abbreviations

IgAV: Immunoglobulin A vasculitis
GC: germinal center
HC: healthy control
PBMC: Peripheral blood mononuclear cell
BCR: B-cell receptor

## Ethics approval and consent to participate

Written informed consents were obtained from parents or guardians of all study participants. The experimental protocol followed the guidelines of the Declaration of Helsinki and was approved by the Human Ethics Committee of Jilin University (Jilin University, Changchun, China).

## Consent for publication

All the authors agreed to publish.

## Availability of data and material

The data used and analyzed in the present study are available from the corresponding author on reasonable request.

## Authors’ contributions

DL and YJ carried out the experiments, and analyzed, and interpreted the data. JW, JL, MX and CL were responsible for the collection of blood samples and the interpretation of the data from the clinical perspective. SY contributed to the conception and design of the study, the analysis and interpretation of the data, and drafting and revising the manuscript. All authors read and approved the final manuscript.

## Acknowledgments

We sincerely thank the patients for their participation in this study, as well as the Key Laboratory of Zoonoses Research of the First Hospital of Jilin University.

